# Miro ubiquitination is critical for efficient damage-induced PINK1/Parkin-mediated mitophagy

**DOI:** 10.1101/414664

**Authors:** Guillermo López-Doménech, Christian Covill-Cooke, Jack H. Howden, Nicol Birsa, Corinne Morfill, Nicholas J. Brandon, Josef T. Kittler

**Affiliations:** Neuroscience, Physiology and Pharmacology, University College London. Gower Street, WC1E 6BT, London, UK.; UCL Institute of Neurology, Queen Square, London, WC1N 3BG, UK.; Neuroscience, IMED Biotech Unit, AstraZeneca, Boston, MA, USA.

**Keywords:** Parkinson’s Disease, Rhot1, Rhot2, ubiquitin, MEFs, giant mitochondria

## Abstract

Clearance of mitochondria following damage is critical for neuronal homeostasis. Here, we investigate the role of Miro proteins in mitochondrial turnover by the PINK1 / Parkin mitochondrial quality control system *in vitro* and *in vivo*. We find that upon mitochondrial damage, Miro is promiscuously ubiquitinated on multiple lysine residues. Combined knockout of both Miro1 and Miro2 or block of Miro ubiquitination and subsequent degradation, lead to slowed mitophagy. In cultured neurons, Miro1 knockout also leads to delayed Parkin translocation onto damaged mitochondria and reduced mitochondrial clearance. *In vivo*, postnatal knockout of Miro1 in hippocampus and cortex disrupts mitophagy and leads to a dramatic age dependent upregulation of the mitofusin mitochondrial fusion machinery. Fluorescence imaging of aged neurons conditionally knocked out for Miro1 and expressing mitoDendra to label mitochondria *in vivo*, reveals that Mfn1 / Mfn2 upregulation leads to enlarged and hyperfused somatic mitochondria. Our results provide new insights into the role of Miro in PINK1/Parkin dependent mitophagy and further suggest that disruption of this regulation may be implicated in human neurological pathology.

## Introduction

Mitochondria are critical for ATP provision and play other essential roles in cells such as buffering calcium and lipid synthesis (MacAskill and Kittler, 2010; Mishra and Chan, 2014; Sheng and Cai, 2012). The tight regulation of mitochondrial transport and distribution is therefore crucial as it enables mitochondria to be delivered and localised to areas where they are needed. The outer mitochondrial membrane (OMM) Miro (mitochondrial Rho) GTPases, Miro1 and Miro2, have emerged as key regulators of mitochondrial trafficking and distribution by both the microtubule and actin cytoskeletons. Miro proteins regulate the activity of the kinesin/dynein motor complexes in coordination with the recruitment and stabilization of myosin-19 to the mitochondrial membrane (Birsa et al., 2013; Fransson et al., 2006; Lopez-Domenech et al., 2018; Stowers et al., 2002). Miro proteins have a C-terminal transmembrane domain for OMM targeting and two GTPase domains flanking two Ca^2+^-sensing EF-hand domains for calcium-dependent mitochondrial stopping (Birsa et al., 2013; Devine et al., 2016; Macaskill et al., 2009; Wang and Schwarz, 2009). Miro proteins may also have other important roles for mitochondrial function as components of mitochondria-ER contact sites and as regulators of mitochondrial calcium homeostasis (Kornmann et al., 2011; Lee et al., 2018; Lee et al., 2016; Niescier et al., 2018).

An accurate mitochondrial quality control system is needed to clear damaged mitochondria that can no longer sustain the cell’s metabolic requirements and that may be a source of damaging reactive oxygen species (ROS) (Covill-Cooke et al., 2018; Pickles et al., 2018). PINK1 (PTEN-induced putative kinase 1; *PARK6*), a mitochondrial serine-threonine kinase and Parkin (*PARK2*), an E3 ubiquitin ligase, are components of a mitochondrial quality control apparatus that promotes the selective turnover of damaged mitochondria through mitochondrial autophagy (mitophagy). PINK1, normally imported into the mitochondrion and cleaved at the inner mitochondrial membrane (IMM), selectively accumulates in its full-length form on the OMM of damaged mitochondria where it phosphorylates the serine 65 (S65) residue of ubiquitin, as well as a conserved residue in the ubiquitin-like domain of Parkin (reviewed in (Harper et al., 2018)). This leads to the recruitment and activation of Parkin from the cytosol to the mitochondria to ubiquitinate various OMM substrates (Cai et al., 2012; Deas et al., 2011; Exner et al., 2012; Lazarou et al., 2015; Sarraf et al., 2013; Vives-Bauza et al., 2010). The ubiquitination of Parkin substrates on the OMM triggers the recruitment of autophagic adaptors (e.g. p62, NDP52, optineurin) and is a crucial step in the clearance of damaged mitochondria through the autophagic pathway (Lazarou et al., 2015; Narendra et al., 2010). Importantly, loss of function mutations in PINK1 and Parkin are associated with rare recessive forms of Parkinson’s Disease (PD) (Thomas and Beal, 2007) supporting an important role for mitophagy in neuronal survival and a link between its dysregulation and neurodegenerative diseases.

Miro proteins are important targets of PINK1/Parkin-dependent ubiquitination and degradation (Birsa et al., 2014; Liu et al., 2012; Sarraf et al., 2013; Wang et al., 2011). Regulation of the Miro trafficking complex by PINK1 and Parkin may serve to block the microtubule based transport and re-distribution of damaged mitochondria, helping to isolate the damaged organelles from the functional mitochondrial network (Wang et al., 2011). PINK1/Parkin-mediated Miro degradation also uncouples mitochondria from actin-dependent trafficking and/or anchorage via rapid loss of Myo19 from the OMM (Lopez-Domenech et al., 2018) suggesting that Miro may also coordinate actin dependent processes during mitophagy. We have previously reported that, in addition to acting as a Parkin substrate, Miro proteins might directly act as receptors for Parkin on the OMM to facilitate Parkin stabilisation (Birsa et al., 2014). Thus, the Miro machinery may play an active role in facilitating mitochondrial Parkin recruitment to damaged mitochondria, a process which may also be regulated by phosphorylation (e.g. by PINK1; (Shlevkov et al., 2016)). Furthermore, Miro degradation was shown to be impaired in fibroblasts and neurons derived from patients with PD (Birsa et al., 2014; Hsieh et al., 2016) although whether Miro is directly involved in the mitophagic clearance of mitochondria and its role in mitochondrial homeostasis *in vivo* remains poorly understood.

Here, we use mouse embryonic fibroblasts (MEFs) knocked out for both Miro proteins, in addition to constitutive knockout (KO) of Miro2 and a conditional mouse knockout (CKO) of Miro1 to investigate their role in mitochondrial turnover by the PINK1/Parkin pathway *in vitro* and *in vivo*. Ubiquitination assays in cells reveal that upon mitochondrial damage Miro is ubiquitinated on multiple lysine residues by Parkin. Blocking Miro ubiquitination stabilises Miro levels upon mitochondrial damage and leads to slowed mitophagy. In addition, loss of both Miro proteins in Miro double-knockout (Miro^DKO^) cells leads to disrupted Parkin translocation and slowed clearance of damaged mitochondria suggesting that a tight temporal regulation of Miro levels at the OMM is required for efficient mitophagy. Deletion of Miro1 in cultured neurons also led to delayed Parkin translocation onto damaged mitochondria and reduced mitochondrial loss. *In vivo*, disruption of mitophagy by postnatal forebrain knockout of Miro1 associates with a dramatic and age dependent upregulation of Mfn1 and Mfn2, which correlates with the re-organisation of the mitochondrial network and pathological mitochondrial hyperfusion. Our results provide new insights into the role of Miro in PINK1/Parkin dependent mitophagy and further suggests that disruption of this regulation may be implicated in neurological damage *in vivo*.

## Results

### Damage induced mitophagy is slowed in Miro^DKO^ cells

Whilst it is well established that Miro proteins are rapidly ubiquitinated by PINK1 and Parkin upon mitochondrial damage (Birsa et al., 2014; Liu et al., 2012)(see also STable1), how important Miro may be for the mitophagic process remains less clear. We previously reported that Miro itself might act as a stabiliser of mitochondrial Parkin acting in the first steps of Parkin-dependent mitophagy (Birsa et al., 2014). To investigate the importance of Miro for damage-induced Parkin stabilisation we used a recently characterised Miro1/2 double knockout cell line (Miro^DKO^ MEFs that lack both Miro1 and Miro2) (Lopez-Domenech et al., 2018). Expressing YFP-Parkin and inducing mitochondrial damage with FCCP is a well-characterised assay for studying the time-dependent translocation of Parkin onto the mitochondria. Using this assay, we followed 4 key stages of Parkin distribution and mitochondrial re-modelling during FCCP induced mitochondrial damage (Fig 1A): i) diffuse Parkin distribution within cells; ii) appearance of sparse Parkin puncta onto mitochondrial units; iii) mitochondrial aggregation and peri-nuclear redistribution of Parkin positive mitochondria; and iv) complete translocation of Parkin across all the mitochondrial network. In a blinded experimental fashion, we were able to compare the above stages of the mitophagic process in Miro^DKO^ cells compared to those found in wild type (WT) control MEFs. At early time-points, 1 and 3 hours of FCCP treatment, we observed a significantly higher proportion of Miro^DKO^ cells with a diffuse distribution of Parkin compared to WT cells (Fig 1B). This correlated with a delay in the appearance of Parkin positive puncta, which peaked at 1 hour of FCCP treatment in WT cells, but which was delayed in Miro^DKO^ cells to between 3 and 6 hours (Fig 1C). In addition, Parkin translocation after mitochondrial damage in WT MEFs induced the remodelling and perinuclear aggregation of mitochondria, with large numbers of cells exhibiting substantial mitochondrial aggregation by 3 hours after damage induction. In contrast, at this time point, this metric was also significantly delayed in Miro^DKO^ MEFs (Fig 1D). Finally, by 6 hours over 50% of WT cells exhibited an almost complete Parkin translocation onto the mitochondrial network, which was greatly reduced in the Miro^DKO^ MEFs (Fig 1E). Therefore, the presence of Miro at the OMM appears to be critical for the efficient recruitment of Parkin to damaged mitochondria.

**Figure 1:**
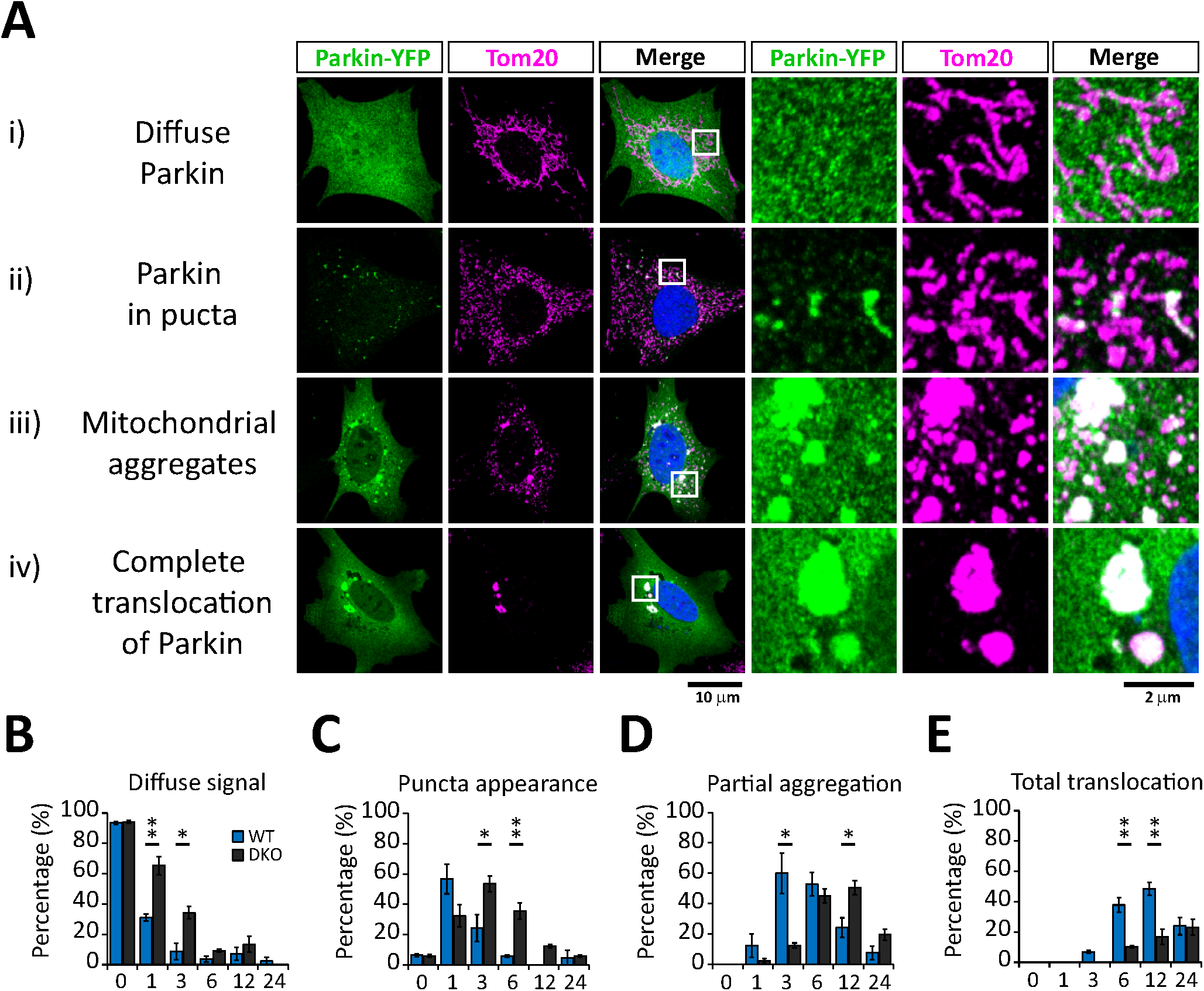
Miro double Knockout cells (Miro^DKO^) have delayed mitophagy upon mitochondrial damage. (A) Representative images (and details from boxed regions) of MEFs cells transfected with YFP-Parkin (green) and treated with FCCP (10 μM) to induce mitophagy. Cells were immunostained with anti-Tom20 (magenta) to reveal the mitochondrial network. Each row (i to iv) represents one stage in the process of mitophagy that has been used to score the progression of mitophagy triggered by a FCCP (10 μM) treatment in WT and Miro^DKO^ cells transfected with YFP-Parkin at different time points (quantification in B to E). From top to bottom: (i and B) “Diffuse Parkin” defines cells in where Parkin expression is homogeously distributed throughout the cytoplasm. (ii and C) Soon after mitochondrial damage Parkin appears enriched in small “puncta” throughout the cytoplasm localizing with isolated mitochondria or portions of mitochondria. (iii and D) These puncta structures soon appears to aggregate in bigger structures accumulating in perinuclear regions. (iv and E) The process of Parkin translocation eventually affects all the mitochondrial population that usually collapses in few large structures possibly forming part of big autolysosomes in the perinuclear region. Error bars represent s.e.m. Significance: *p<0.05 and **p<0.01

### Miro is promiscuously ubiquitinated on multiple lysine residues

To further understand the role of Miro in mitophagy and mitochondrial clearance we set out to further characterise lysine residues important for Miro ubiquitination in cells. Several lysine residues have been identified as sites for Miro ubiquitination, both from *in vitro* studies (Kazlauskaite et al., 2014) and from unbiased mass spectrometric analysis of ubiquitination in cells (STable1). We generated a series of single point mutants in Miro1 to some of these sites along with a compound mutation lacking five key potential sites (Miro1^5R^ hereafter). We selected the residues K153, K187 and K572 because they were identified in several studies as being ubiquitinated upon mitochondrial damage, and K182 and K194 due to their close proximity to K187, likely a critical ubiquitination target along with K572 (Fig 2A and STable1). In addition, we synthesized a Miro1 mutant construct in which all 39 lysine residues of Miro1 were replaced by arginine (Miro1^allR^). Because lysine 572 (K572) was previously demonstrated to be the main site for Parkin dependent ubiquitination *in vitro* (Kazlauskaite et al., 2014; Klosowiak et al., 2016) we also generated an additional construct where this residue was reintroduced on the Miro1^allR^ backbone (Miro1^R^572^K^). All constructs were efficiently expressed and effectively targeted to mitochondria in Miro^DKO^ MEFs (Fig 2B). We found that mutation of any one site (including K572) did not significantly alter Miro1 expression levels (Fig 2C). We then tested the ability of these constructs to be ubiquitinated upon their expression in SH-SY5Y cells stably overexpressing ^Flag^Parkin, where robust damage-induced Miro ubiquitination has been previously reported (Birsa et al., 2014). Surprisingly, the compound mutation of 5 of the key potential sites (including K572) of Miro ubiquitination in the Miro1^5R^ construct led to an only partial decrease in the levels of damage-induced Miro ubiquitination (Fig 2D and E) suggesting that in cells, as opposed to *in vitro*, multiple lysine residues may serve as substrates for Miro ubiquitination and not just K572. Only by mutating all potential Miro1 ubiquitination sites could we completely block its damage-induced ubiquitination (Fig 2D and E), thus Parkin exhibits significant promiscuity in targeting lysine residues on Miro1. In agreement with the conclusion that no individual site is necessary and sufficient to rescue Miro1 ubiquitination, reintroducing K572 on the Miro1^allR^ background was unable to rescue the extent of damage-induced Miro1 ubiquitination or degradation (Fig 2D, E and G). We then expressed wild type Miro1, Miro1^5R^ or Miro1^allR^ and investigated the impact on time dependent FCCP induced Miro1 loss. Blocking all Miro1 ubiquitination with the Miro1^allR^ construct correlated with the stabilisation of Miro1 protein levels at both the 3 hour and 6 hour time points which does not occur when Miro1 ubiquitination is only reduced in the Miro1^5R^ condition (Fig 2H). Unexpectedly, however, we also noted that the damage-induced loss of PDH-E1α (a mitochondrial matrix protein) was reduced in both Miro1^5R^ and Miro1^allR^ expressing cells (Fig 2I), while Parkin levels also appeared to be stabilised (Fig 2F and S1). This suggests that reducing Miro1 ubiquitination may impact on the turnover of damaged mitochondria and Miro ubiquitination may not just be required to stop the trafficking of damaged mitochondria but may be directly involved in facilitating the mitophagic process.

**Figure 2:**
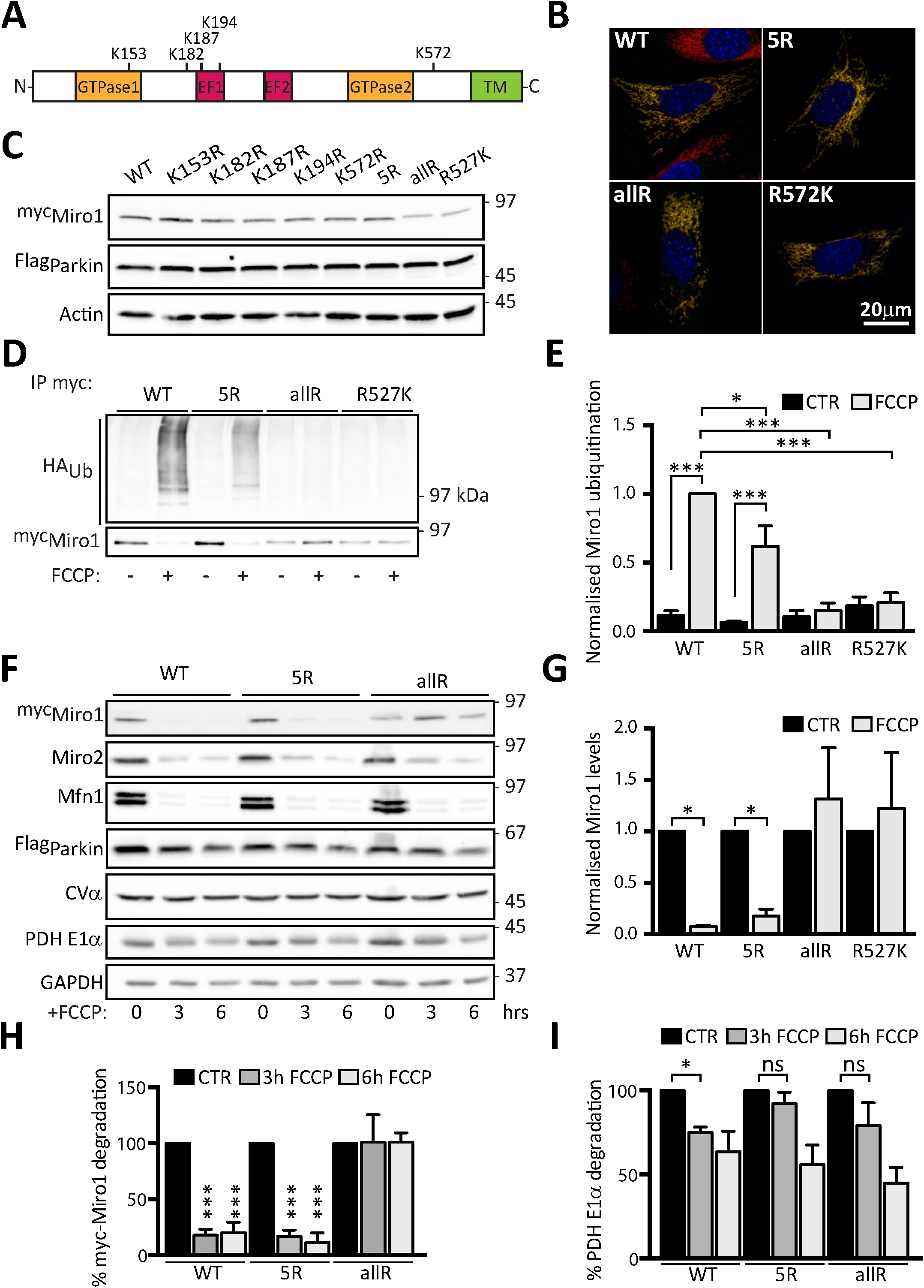
Miro1 ubiquitination is a required early step during the mitophagic process. (A) Schematic representation of Miro1 domains highlighting key lysine (K) residues reported to be ubiquitinated upon mitochondria damage. (B) Representative images showing full length control and selected Miro1 Lysine (Lys)-mutants (5R-myc, allR-myc and R572K-myc) expressing in Miro^DKO^ MEF cells. All constructs (myc-tag, green) localise to mitochondria (Tom20, red). (C) Immunoblot showing expression of Miro1 Lys-mutants in Flag-Parkin expressing SH-SY5Y cells. (D) Ubiquitination assay showing that Miro1 Lys-mutants have reduced ubiquitination upon FCCP treatment (1h, 10 μM) in Flag-Parkin overexpressing SH-SY5Y. The effect is quantified in (E). N=4 for WT and 5R and n=3 for allR and R572K, ANOVA test with Sidak’s multiple comparison post hoc test. (F) Representative western blot showing a degradation assay in Flag-Parkin overexpressing SH-SY5Y cells transfected with WT, 5R or allR Miro1 contructs and treated with FCCP (10 μM) for 3 or 6 hours. (G) Quantification of WT, 5R, allR and R572K Miro1 degradation in (D) N=4, ANOVA test with Tukey’s multiple comparison post hoc test. (H) Quantification of WT, 5R and allR Miro1 degradation in (F). (I) Quantification of the matrix protein PDH E1α in (F). N=4, ANOVA test with Sidak’s multiple comparison post hoc test. Error bars represent s.e.m. Significance: *p<0.05, **p<0.01 and ***p<0.001.

### Ubiquitination of Miro1 is a required step during mitophagy

To further investigate the importance of Miro1 ubiquitination for Parkin translocation and mitochondrial clearance we performed rescue experiments in Miro^DKO^ MEFs by re-expressing either wild-type Miro1 or the ubiquitination mutant, Miro1^allR^. All key stages of damage induced Parkin re-distribution and mitochondrial re-modelling including loss of diffuse Parkin within cells, appearance of Parkin puncta, mitochondrial aggregation and peri-nuclear redistribution and the complete translocation of Parkin on mitochondria were examined and found to be significantly reduced in cells expressing Miro1^allR^ compared to WT Miro1 (Fig 3A-E) suggesting that mitophagy was slowed. Indeed, when we quantified mitochondrial loss as a read out of the efficiency of mitochondrial clearance we observed a dramatic reduction of damage induced mitochondrial turnover in mutant Miro1^allR^ expressing cells (Fig 3A and F). This suggests that Miro accomplishes two differentiated roles in regulating mitophagy. One, allowing the recruitment of Parkin to the mitochondria to initiate the ubiquitination of targets, and a second one that requires the ubiquitination of Miro that leads to the stabilization of Parkin in the OMM and ultimately the remodelling and degradation of mitochondria. Thus, the presence of Miro proteins and their ubiquitination appear to be equally important for ensuring the efficient stabilisation of Parkin on the mitochondrial membrane during damage and for the subsequent clearance of damaged mitochondria.

**Figure 3:**
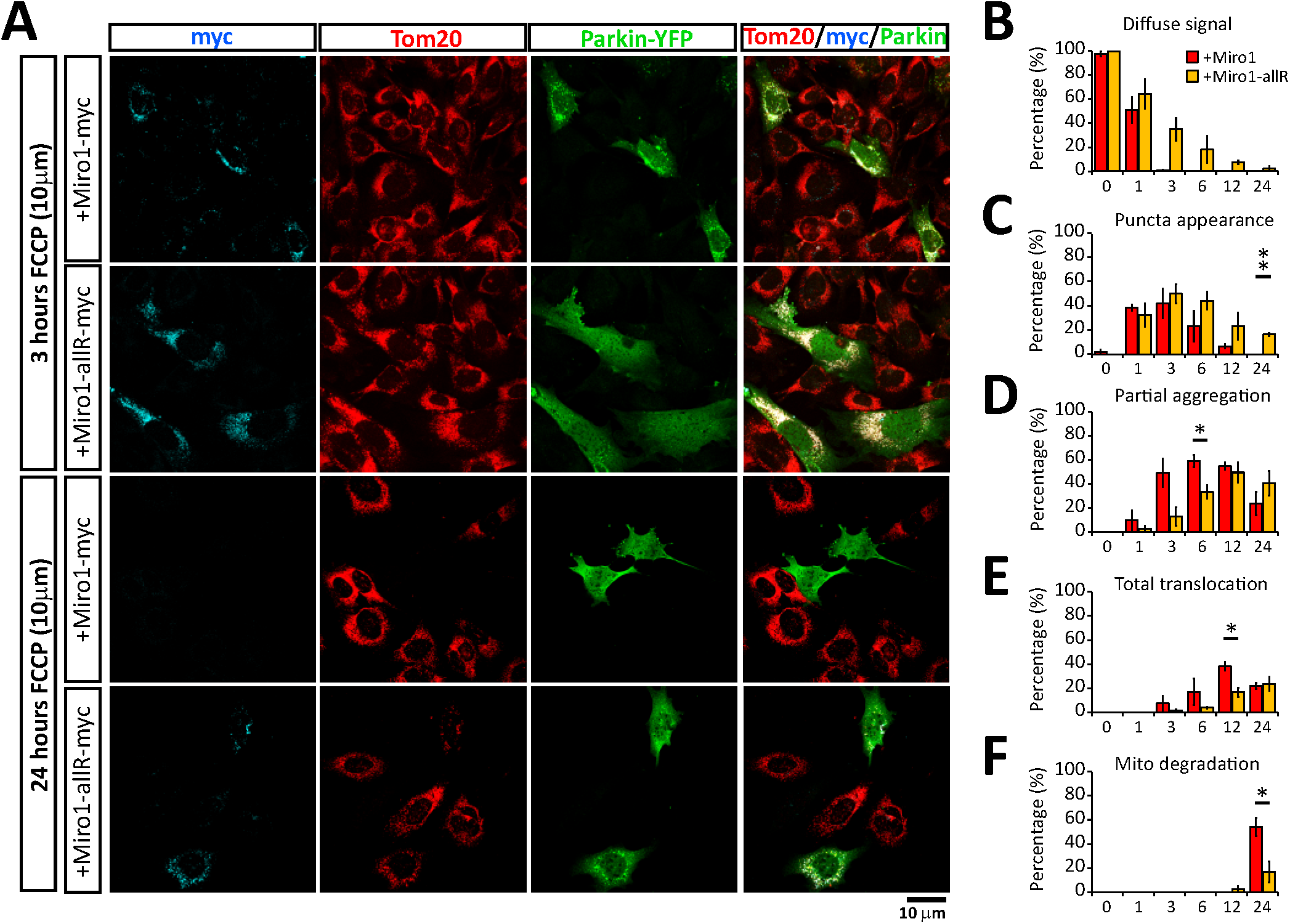
Miro ubiquitination is required for mitochondrial degradation through mitophagy. (A) Representative images at 3 and 24 hours of FCCP treatment (10 μM) of YFP-Parkin (green) expressing Miro^DKO^ cells co-transfected with full length Miro1-myc or the Miro1^allR^ construct (cyan). Tom20 (red) was used to reveal the mitochondrial network. (B - E) Shows the quantification of the mitophagic process scored as: (A) Diffuse Parkin; (B) Parkin puncta; (D) Mitochondrial aggregates and (E) Complete Parkin translocation. (F) Quantification of the mitochondrial clearance defined as cells with the complete absence of a mitochondrial compartment. Error bars represent s.e.m. Significance: *p<0.05 and **p<0.01

### Miro1 is critically important in primary neurons to recruit Parkin to mitochondria to trigger mitophagy

Although Miro^DKO^ MEFs provide a powerful system to study the Miro-dependency of mitophagy, neurons represent a more physiologically and pathologically relevant model. A number of previous studies have investigated conditions to induce Parkin translocation in neurons with some debate with respect to the optimal conditions and timescales that enable Parkin translocation without significant cytotoxicity (Cai et al., 2012; Joselin et al., 2012; Van Laar et al., 2011; Wang et al., 2011). In fact, caspase inhibitors have often been required to study Parkin-mediated mitophagy at longer timescales (Cai et al., 2012; Lazarou et al., 2015). Therefore, we initially investigated conditions favourable for driving Parkin translocation in ^YFP^Parkin transfected mouse primary cultured neurons, without leading to significant cell death. In our hands, treatment with commonly used concentrations of FCCP or antimycin-A led to a significant amount of cell death within 5 hours, close to the levels of death induced by an excitotoxic glutamate insult (Fig S2A and B), precluding using these conditions to accurately study Parkin translocation in neurons. The potassium ionophore valinomycin depolarizes the mitochondria without affecting the pH gradient leading to a strong PINK1 activation (Kondapalli et al., 2012). In contrast to FCCP or antimycin-A, 1μM valinomycin treatment in primary mouse neurons initially led to a rapid remodelling of the mitochondrial network, followed by the induction of Parkin translocation (Fig 4A) without sustaining significant neuronal death (Fig S2A and B). Initially, Parkin accumulated in hot spots on the mitochondria, which eventually became Parkin rings entirely surrounding mitochondria, as could be clearly observed in deconvoluted confocal images, a method that increases the resolution of images by reversing the blurring caused by the limited aperture of the objective (Fig 4B). Using the same conditions, we then investigated Parkin translocation upon knockout of Miro1. In contrast to WT neurons, we observed a dramatic delay in Parkin translocation in Miro1^KO^ neurons (Fig 4A and B), as could be observed by reduced overlap between Parkin and the mitochondrial marker mtDsRed in line scans and in the extent of co-localisation between ^YFP^Parkin and mtDsRed (Fig 4C, D and E). Subsequent to remodelling and Parkin translocation, we could also observe a loss of somatic mitochondrial content as mitochondria begin to be cleared by the mitophagic process, which was also significantly reduced in Miro1^KO^ neurons (Fig 4F-H). Thus, Miro1 appears to be critically important in primary neurons for the stabilisation of Parkin on the OMM upon mitochondrial damage, which is a critical step for damage-induced mitochondrial clearance.

**Figure 4:**
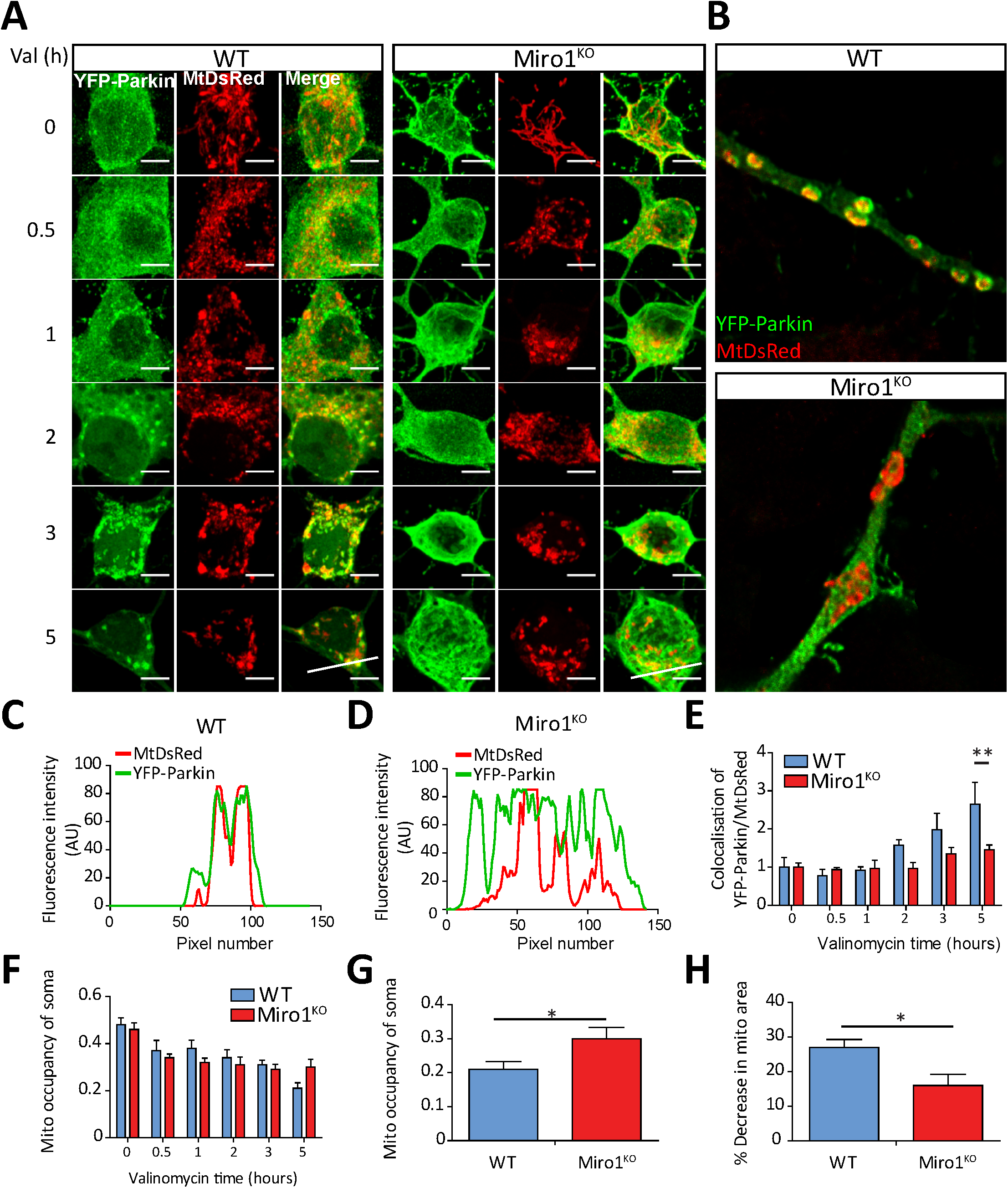
Loss of Miro1 delays Parkin recruitment to damaged mitochondria in neurons. (A) Representative confocal images of the soma of WT and Miro1^KO^ neurons after valinomycin treatment (Scale bars = 5µM). (B) Deconvoluted confocal images of Parkin recruitment after 5 hours of valinomycin treatment in WT and Miro1^KO^ neurons. (C) Fluorescent linescans of YFP-Parkin and MtDsRed signal in WT and Miro1^KO^ neurons after 5 hours of valinomycin treatment (Lines shown in A). (D) Quantification of parkin recruitment (YFP-parkin signal overlapping MtDsRed signal – YFP-Parkin signal not overlapping MtDsRed signal) in WT and Miro1^KO^ neurons following valinomycin treatment (** - P<0.01 two-way ANOVA, n≥15 cells all conditions). (E) Quantification of mitochondrial occupancy (Area of MtDsRed signal in soma / Entire area of soma) in the soma of WT and Miro1^KO^ neurons following valinomycin treatment. (F) Mitochondrial occupancy in the soma of WT and Miro1^KO^ neurons following 5 hours of valinomycin treatment (* - P<0.05, Unpaired t-test). (G) Quantification of mitochondrial clearance from the soma of WT and Miro1^KO^ neurons following 5 hours of valinomycin treatment (* - P<0.05, Unpaired t-test). Error bars represent s.e.m. Significance: *p<0.05 and **p<0.01

### Deletion of Miro1 leads to an upregulation of mitofusins and the mitophagic machinery *in vivo*

Given that defects in mitophagy can lead to an age-dependent accumulation of mitochondrial dysfunction, we next sought to investigate how altered mitophagy through loss of Miro1 or Miro2 would affect mitochondrial homeostasis and the expression levels of key mitochondrial proteins in mouse brain, where neurons can develop on an extended timescale compared to cell cultures. Constitutive loss of Miro1 is lethal postnatally (Lopez-Domenech et al., 2016; Nguyen et al., 2014), unlike Miro2, in which global knockout has no effect on viability (Lopez-Domenech et al., 2016). We therefore used a conditional Miro1 model where Miro1-floxed mice were crossed with mice expressing Cre-recombinase under the CaMKIIα promoter, allowing the impact on mitochondrial homeostasis of knockout of Miro1 in postnatal excitatory neurons (Miro1^CKO^ from hereon) to be compared to wild type neurons. We also analysed the impact of knocking out Miro2, which can also be a substrate for damage-induced ubiquitination (Birsa et al., 2014; Ordureau et al., 2018; Sarraf et al., 2013) and might help shed light into specific roles of Miro1 and Miro2 in regulating mitochondrial homeostasis *in vivo*. Hippocampal lysates were then prepared at 4 and 12 months of age from WT, Miro1^CKO^ and Miro2 constitutive knockout (Miro2^KO^) animals, and studied by western blotting. At 4 months of age, no difference in Parkin and PINK1 levels were observed between WT, Miro1^CKO^ and Miro2^KO^ animals (Fig 5A and C). To test for variations in the activity of Parkin-mediated mitophagy *in vivo* the levels of Mfn1, Mfn2 and VDAC1 - well characterized Parkin substrates - were also probed (Chen and Dorn, 2013; Gegg et al., 2010; Geisler et al., 2010; Sarraf et al., 2013). Interestingly, at 4 months, an increase in Mfn1 was observed in Miro1^CKO^ in comparison to WT animals, though no significant increase in VDAC1 was observed (Fig 5A and C). Loss of Miro1 or Miro2 had no effect on basal mitochondrial content at either 4 or 12 months, as observed by no changes in ATP5a (the alpha subunit of the ATPase in complex V of the electron transport chain) (Fig 5B and D). Strikingly, however, the levels of both Mfn1 and Mfn2 in the Miro1^CKO^ at 12 months were substantially and consistently increased in comparison to WT hippocampal lysates; an effect that was not observed in the Miro2^KO^ mice (Fig 5B and D). To confirm the increase in Mfn1 and Mfn2 protein levels, coronal brain slices from 12 months animals were stained. Indeed, a large increase in both Mfn1 and Mfn2 staining was observed in principal neurons of the cortex from the Miro1^CKO^ brains in comparison to WT brains (Figure 5E). Alongside the increase in Mfn1 and Mfn2 in 12 month old Miro1^CKO^ mice, our western blot analysis revealed that there is also a significant increase in PINK1 levels and the appearance of an upper band of Parkin, suspected of being an auto-ubiquitinated form of Parkin (Figure 5B) (Chaugule et al., 2011; Wauer and Komander, 2013). As Miro1 is knocked out in only principal cells, it is likely that the changes observed in the 12 month Miro1^CKO^ tissue lysates are in fact much higher than the western blot data indicate. In summary, loss of Miro1 but not Miro2 leads to an upregulation of the mitophagic machinery in an age-dependent manner.

**Figure 5:**
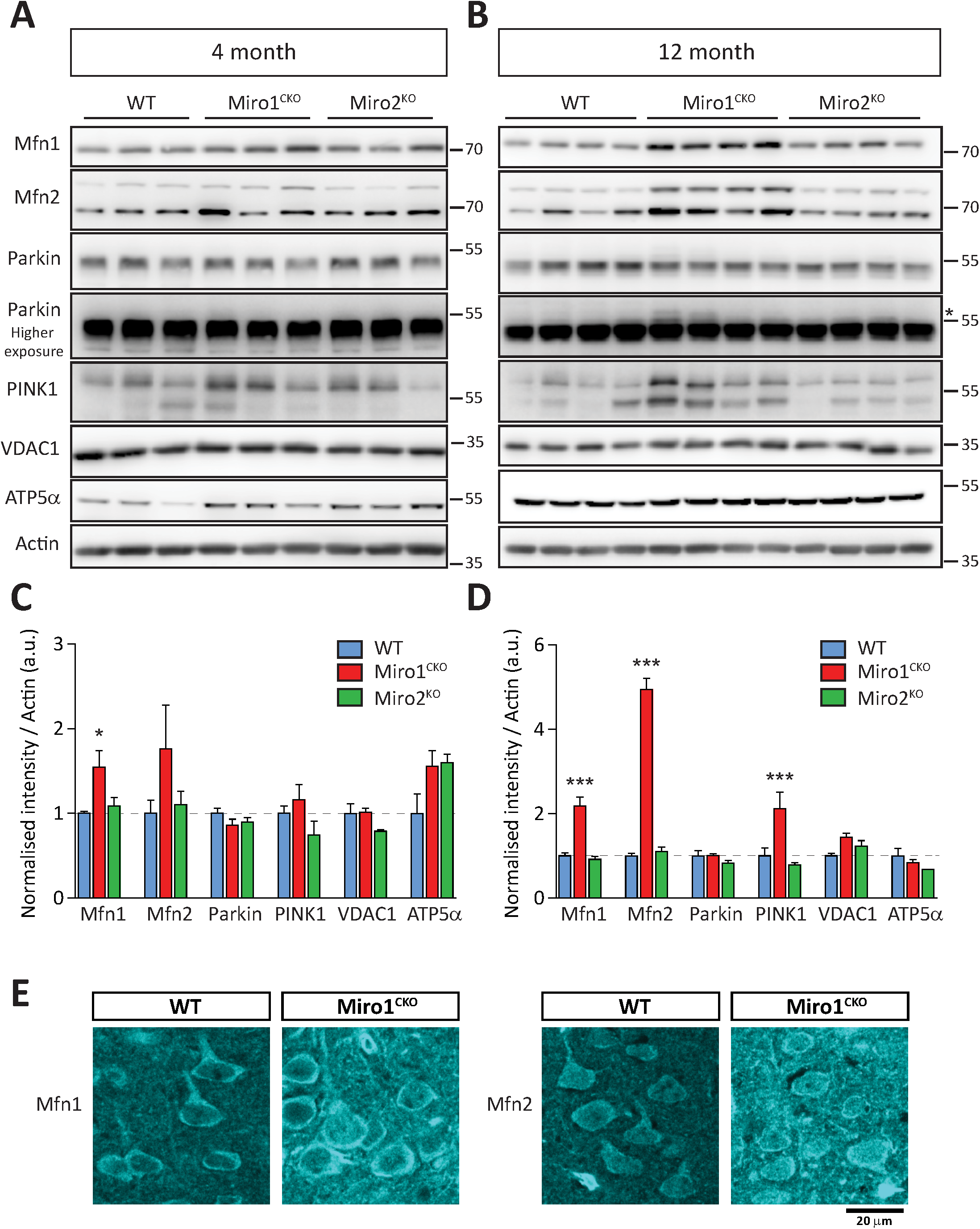
Loss of Miro1, but not Miro2, leads to an age-dependent increase in mitophagy machinery. (A) Representative blots of 4 month hippocampal lysates from three mice of each genotype (WT, Miro1^CKO^ and Miro2^KO^). (B) Representative blots of 12 month hippocampal lysates from four mice of each genotype (WT, Miro1^CKO^ and Miro2^KO^). The asterisk (*) indicates a possible auto-ubiquitinated form of Parkin. (C and D) Quantification of protein levels in 4 month lysates (n=3 animals/genotype) and 12 month lysates (n=4 animals/genotype), respectively. (E) Representative images of Mfn1 and Mfn2 staining in cortical slices from 12 month WT and Miro1^CKO^ mice. All data represent mean ± SEM. * and *** represent p<0.05 and p<0.001, respectively, using a one-way ANOVA with post-hoc Dunnett’s test. Error bars represent s.e.m. Significance: *p<0.05, **p<0.01 and ***p<0.001.

### Loss of Miro1 in principal neurons induces the appearance of giant mitochondria with altered morphology

The dramatic upregulation of Mfn1 and Mfn2 in Miro1^CKO^ neurons suggests a pathological re-configuration of the mitochondrial network due to an altered balance of the fission/fusion dynamics caused by disruption of PINK1/Parkin-mediated mitophagy. To further address this *in vivo*, we crossed the Miro1^CKO^ mouse line with a mouse line that allows conditional Cre-recombinase-dependent expression of the mitochondrial reporter mitoDendra (Fig 6A). Crossing the mitoDendra line with Cre recombinase led to robust expression of mitoDendra in CamKIIα driven Cre expressing cells, as observed by significant mitoDendra expression in the forebrain and hippocampus (Fig 6B). This approach allowed us to visualise the mitochondrial network, specifically in principal neurons of cortex and hippocampus from aged animals. In control mice (heterozygous for the Miro1 conditional allele - Miro1(Δ/+)) mitoDendra labelled mitochondria appeared as an elongated network of various sized mitochondrial elements. Using a similar approach, we also generated aged matched Miro2^KO^ mice with mitochondria labelled with mitoDendra in Cre-expressing cells (Fig 6C) which showed similar features as in control animals (Fig 6C). In stark contrast, mitoDendra labelled mitochondria in Miro1^CKO^ cells at 1 year exhibited a dramatic remodelling of somatic mitochondria (Fig 6C and S4). This was particularly evident in the cortex where labelling was sufficiently sparse to easily identify pyramidal cell bodies. Zooms of the mitochondrial network in Miro1^CKO^ cell bodies revealed large mitochondrial units of accumulated and hyperfused “giant” mitochondria (Fig 6C), which were further resolved using high resolution imaging (Airyscan - Zeiss), a method that increases imaging resolution by a factor of 1.7 (Fig S4). Giant mitochondria have been identified in a number of models where either mitochondrial fission / fusion proteins or mitophagy has been altered (Chen et al., 2007; El Fissi et al., 2018; Kageyama et al., 2014; Kageyama et al., 2012; Yamada et al., 2018a; Yamada et al., 2018b). Interestingly, we observed that a proportion of Miro1^CKO^ cells presenting giant mitochondria showed an accumulation of ubiquitin surrounding the aberrant mitochondrial particles similar to what happens during ageing or Lewy Body Disease (Fig 6D) (Hou et al., 2018). Finally, to address the impact on the mitochondrial network with higher resolution we performed electron microscopic analysis from control or Miro1^CKO^ tissues. While mitochondria in control neurons from 4 months old animals appeared with the expected elongated morphology (Fig 6E) we observed that mitochondria in Miro1^CKO^ cells presented increased size and altered morphology (Fig 6E). The mitochondrial units appeared swollen and rounded and the intra-mitochondrial space less electron dense than that from control mitochondria (Fig 6E). Importantly, this effect was greatly exacerbated in neurons from 12 months old animals (Fig 6E) indicating that alterations in mitochondrial morphology due to loss of Miro1 are progressive, and presumably in response to accumulation of Mfn1 and Mfn2 due to a defective mitochondrial clearance.

**Figure 6:**
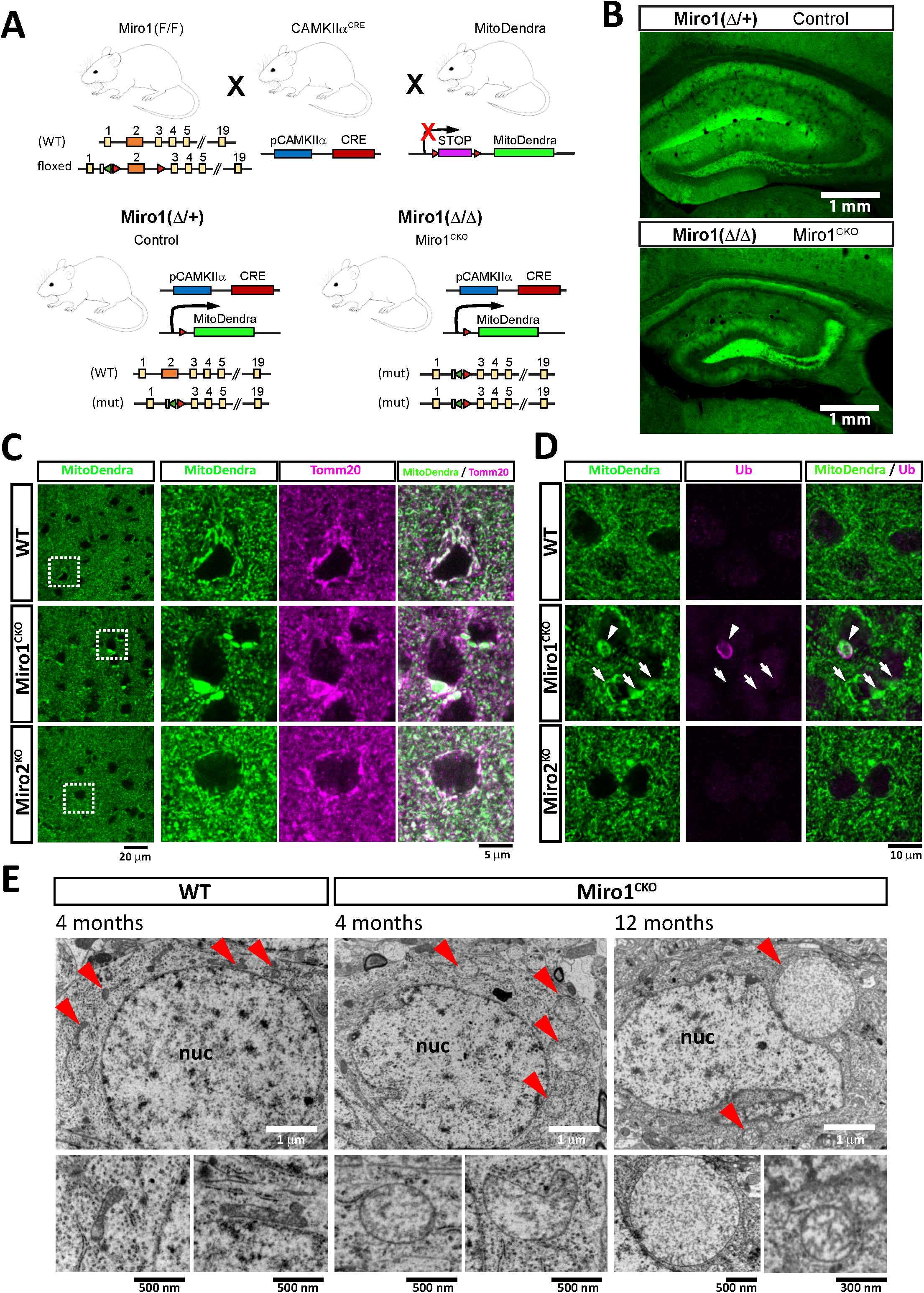
Loss of Miro1 in mature neurons associates with age-dependent accumulation of defective mitochondria in neurons. (A) Schematic depicting the mouse lines used to generated control (Miro1(Δ/+)) and conditional knockout animals (Miro1(Δ/Δ)) with labelled mitochondria in only the cells of interest. (B) representative images of hippocampal regions from Control and Miro1^CKO^ crossed with MitoDendra animals. (C) Details from cortical regions of WT, Miro1^CKO^ and Miro2^KO^ animals at 12 months of age showing large mitochondrial structures occurring only in the case of Miro1^CKO^ animals (arrows). These large mitochondria present enhanced ubiquitin signaling (arrowhead) in a subset of cells (D). (E) Electronic microcopy images of WT and Miro1^CKO^ brains showing the ultrastructure of the mitochondrial compartment. Red arrows point to mitochondria in low magnification images. The mitochondrial units in Miro1^CKO^ cells are bigger compared to mitochondria from WT cells at 4 months of age. This process is progressive as mitochondrial structures appear importantly enlarged and with reduced electrondensity inside the mitochondria at 12 months of age.

## Discussion

Here, we characterise the role of Miro proteins in damage-induced PINK1/Parkin-dependent mitophagy using Miro1/2 DKO mouse embryonic fibroblasts, primary Miro1 constitutive knockout mouse hippocampal neurons and conditional Miro1 knockout neurons *in vivo*.

We previously showed that Miro1 promotes the recruitment and stabilisation of mutant forms of Parkin (characterised by delayed translocation) onto damaged mitochondria in HEK cells suggesting that Miro may play a role in stabilising Parkin on the OMM (Birsa et al., 2014). This would suggest that Parkin binding by Miro, in conjunction with Parkin activation by phospho-ubiquitin, could act together to stabilize Parkin on the OMM, providing a mechanism to tune Parkin levels to that of substrates on the OMM. In agreement with a key role for Miro in stabilising mitochondrial Parkin, we now show that deleting all Miro in Miro^DKO^ cells leads to dramatically reduced Parkin translocation and accumulation onto mitochondria upon damage. Thus, our results further support our previous proposal that Miro forms part of a Parkin receptor complex on the OMM important for tuning Parkin-mediated mitochondrial quality control. This tuning may itself be further regulated by Miro phosphorylation (Shlevkov et al., 2016). Intriguingly however, Miro ubiquitination also appears to be required to allow the later stages of mitophagy to progress. The slowed Parkin translocation upon expression of the Miro^allR^ may be because Miro ubiquitination is specifically required for Parkin stabilisation and amplification of the ubiquitin signal. Indeed, we previously showed that the pool of ubiquitinated Miro remains localized on the OMM for some considerable time prior to it being turned over, dependent on the activity of the proteasome (Birsa et al., 2014). Thus, Miro ubiquitination (rather than ubiquitination-induced Miro degradation) may directly act as a rapid signal on the OMM for further Parkin stabilisation and amplification of the mitophagic process. A number of mechanisms could underpin this. For example, ubiquitinated Miro could, in addition to stabilising Parkin, be key for the recruitment of other factors such as autophagy receptors to the mitochondrial OMM to facilitate the clustering of damaged mitochondria prior to their clearance by mitophagy.

We also provide new insight into the regulation of Miro turnover and ubiquitination dynamics by PINK1/Parkin. A previous report analysing Miro ubiquitination demonstrated robust mono-ubiquitination of Miro on residue K572 by Parkin *in vitro* (Klosowiak et al., 2016). In contrast, global Mass spectrometric analysis of substrate ubiquitination sites in cells revealed a number of sites upon damage including K107, K153, K187, K194, K230, K235, K249, K330, K512 and K572 on Miro1 and K119, K164, K214 and K410 on Miro2 (Boeing et al., 2016; Kazlauskaite et al., 2014; Ordureau et al., 2015; Ordureau et al., 2014; Povlsen et al., 2012; Sarraf et al., 2013; Wu et al., 2015). Finally, several other reports looking at steady ubiquitination levels revealed further additional sites (Kim et al., 2011; Lumpkin et al., 2017; Mertins et al., 2013; Udeshi et al., 2013; Wagner et al., 2011; Wagner et al., 2012). In our experiments, we did not find that the Miro1 K572 ubiquitination site alone was sufficient for mediating Miro1 ubiquitination and turnover, even in dopaminergic SH-SY5Y neuroblastoma cells stably expressing Parkin, which show high levels of damage induced Parkin activity. Although Miro1 K572 ubiquitination by Parkin may be important, and could, for example, impact the rate or kinetics of Miro ubiquitination by Parkin, importantly, we did not find any substantial difference in the ubiquitination kinetics between Miro^R^572^K^ (where K572 is the only lysine replaced in the Miro1^allR^) and Miro1^allR^. Thus, ubiquitination of Miro1 by Parkin requires the ubiquitination of multiple lysine residues to regulate Miro stability. However, our current data does not preclude the possibility that ubiquitination of K572 acts to prime subsequent ubiquitination of other sites. In fact, it is possible that Miro1 ubiquitination at K572 will occur in a cell-specific manner or have a cell-specific impact on Miro ubiquitination kinetics, depending on the expression levels of Parkin or of other regulatory factors such as Miro phosphorylation state and Miro protein kinases or phosphatases (Shlevkov et al., 2016).

We previously reported that Miro proteins interact with Myo19 and are critical for regulating Myo19 recruitment and stability on the OMM, and that Miro loss upon mitochondrial damage leads to a rapid loss of Myo19 from the outer mitochondrial membrane. PINK1/Parkin-mediated Miro ubiquitination and degradation may therefore also be important for the uncoupling of mitochondria from an actin-dependent trafficking and/or anchorage via Myo19 degradation, perhaps to facilitate other actin-dependent mechanisms such as recruitment of the Myosin VI motor (Kruppa et al., 2018). Thus, uncoupling mitochondria from the actin cytoskeleton via myosin degradation may be an important step in early stages of the mitophagic process or an additional subsequent checkpoint during mitophagy dependent on Miro ubiquitination and/or removal.

Using constitutive and conditional mouse knockout strategies we also demonstrate that Miro1 is important for mitophagy in neurons, both *in vitro* and *in vivo*. Using the mitochondrial uncoupler valinomycin, we established conditions to allow for robust Parkin translocation without causing the significant neuronal death observed with high doses of other commonly used inducers of mitophagy such as FCCP or antimycin-A. In accordance to an earlier report, we found Parkin translocation onto mitochondria to be considerably slower than in non-neuronal cells (Cai et al., 2012). Miro1^KO^ neurons exhibited dramatically reduced levels of Parkin translocation onto mitochondria in cell soma, dendrites and axons compared to wild type neurons. This also correlated with reduced mitochondrial clearance at later time points.

Importantly, we also identified hallmarks of altered mitochondrial quality control *in vivo*. Constitutive Miro1 deletion leads to perinatal lethality and disrupted neuronal development (Lopez-Domenech et al., 2016; Nguyen et al., 2014). To avoid this complication we looked in brains of conditional CaMKIIα-Cre-driven postnatal deletion of Miro1 in hippocampus and cortex. Probing lysates from Miro1^CKO^ animals in adult and aged mouse brains (1 year) revealed a dramatic increase in the expression levels of Mfn1 and Mfn2, which was also already evident at 4 months of age for Mfn1. Alterations in Mfn1 and Mfn2 have been observed in several models of mitophagic dysfunction and neurodegeneration (Chen and Dorn, 2013; Gong et al., 2015; Rocha et al., 2018; Wang et al., 2015). Despite the large increases in Mfn1 and Mfn2 at 12 months, no increase in VDAC1 was observed. Though VDAC1 is a substrate for Parkin it is dispensable for the mitophagic loss of mitochondria (Narendra et al., 2010) which may therefore, explain the disparity in the changes of different Parkin substrates in the Miro1^CKO^ mice. Upregulation of the mitochondrial fusion machinery and in particular gain of function mutations of Mfn2 associated with Charcot Marie Tooth-2 (CMT2) pathology are linked to mitochondrial remodelling and the accumulation of hyperfused giant mitochondria in the neuronal somas (El Fissi et al., 2018; Santel and Fuller, 2001). To facilitate determining the impact of Mfn upregulation *in vivo* we generated a new mouse line where mitoDendra is expressed solely in neurons expressing CamKIIα-Cre driven Miro1 recombination allowing identification of Miro1 deleted neurons. Conventional confocal or high resolution (Airyscan) imaging of the mitoDendra labelled mitochondrial network in Miro1 deleted neurons at 1 year of age revealed a dramatic hyperfusion of the mitochondrial network and large mitochondrial structures located in the cell bodies. This is similar to previous reports of collapsed mitochondrial structures identified in a number of models where either mitochondrial fission / fusion proteins or mitophagy has been altered (Chen et al., 2007; El Fissi et al., 2018; Kageyama et al., 2014; Kageyama et al., 2012; Yamada et al., 2018a; Yamada et al., 2018b). Alongside the increase in Mfn1 and Mfn2 in 12 month old Miro1^CKO^ mice we also observed a significant increase in PINK1 levels and the appearance of an upper band of Parkin, suspected of being an auto-ubiquitinated form of Parkin (Figure 5B) (Chaugule et al., 2011; Wauer and Komander, 2013). Parkin auto-ubiquitination leads to the activation of Parkin E3-ligase activity, with sustained auto-ubiquitination being proposed to be indicative of a pathological state of Parkin activation (Chaugule et al., 2011). Furthermore, overexpression of PINK1 is known to cause mitochondrial dysfunction (Yang et al., 2008) by promoting excessive mitochondrial fusion. It is unclear if these alterations in Miro1^CKO^ mouse are due to an overactive state of both Parkin modification and PINK1 overexpression or rather they respond to a compensatory mechanism to counterbalance dysfunctional mitophagy.

Our findings provide new insights into the requirement of Miro1 and Miro2 for mitophagy following mitochondrial damage and highlight its importance for mitochondrial homeostasis *in vitro* and *in vivo*. Moreover, we uncover a role of Miro as part of the Parkin receptor complex on the OMM and its potential role in tuning Parkin-mediated mitochondrial quality control. This may open new strategies to target mitochondrial dysfunction in PD pathogenesis and other related diseases with alterations in the clearance of dysfunctional mitochondria.

## Acknowledgments

We thank all members of the Kittler lab for helpful discussions and comments on the manuscript. This work has been funded by an ERC starting grant (282430), research prize from Lister Institute for Preventive Medicine and grants from Medical Research Council to J. T. K.. G. L-D. is supported by a AstraZeneca postdoctoral fellowship. C.C-C. is in the MRC LMCB PhD programme at UCL (MRC studentship 1368635). J. H. is the recipient of a BBSRC CASE Award (BB/P504865/1) Ph.D with AstraZeneca.

## Materials and Methods

### Animals

The *Rhot1* (MBTN_ EPD0066_2_F01; Allele: *Rhot1*^*tm1a(EUCOMM)Wtsi*^) and *Rhot2* (MCSF_ EPD0389_5_A05; *Rhot2*^*tm1(KOMP)Wtsi*^) mice lines were obtained from the Wellcome Trust Sanger Institute as part of the International Knockout Mouse Consortium (IKMC) (Skarnes et al., 2011) and previously published (Lopez-Domenech et al., 2016). CAMKIIα-CRE strain has been described previously (Mantamadiotis et al., 2002). MitoDendra (B6;129S-Gt(ROSA)26Sor^tm1(CAG-COX^8^a/Dendra^2^)Dcc^/J) line was obtained from The Jackson Laboratory and was previously described (Pham et al., 2012). Animals were maintained under controlled conditions (temperature 20 ± 2°C; 12 hour light-dark cycle). Food and water were provided *ad libitum*. All experimental procedures were carried out in accordance with institutional animal welfare guidelines and licensed by the UK Home Office in accordance with the Animals (Scientific Procedures) Act 1986. All data involving procedures carried out on in animals is reported in compliance with ARRIVE guidelines (Kilkenny et al., 2010)

### DNA constructs

cDNA construct encoding mycMiro1 has been previously described (Macaskill et al., 2009). K153R, K182R, K187R, K194R, K572R and 5R mycMiro1 (K153R, K182R, K187R, K194R and K572R) were made by site-directed mutagenesis on the mycMiro1 backbone. The Miro1^allR^ construct was purchased from Life Technologies and was inserted into the pML5-myc backbone by restriction digestion. R572K mycMiro1 was made by site-directed mutagenesis on the allR mycMiro1 backbone (and has all K residues mutated to R apart from K572). pRK5-HA-Ubiquitin was from Addgene [plasmid #17608, (Lim et al., 2005)]. YFP-Parkin was from Addgene [plasmid #23955, (Narendra et al., 2008)]. MtDsRed was previously described (Macaskill et al., 2009).

### Antibodies

Primary antibodies for western blotting: mouse anti-Mfn1 (Abcam ab57602, 1:500), mouse anti-Mfn2 (NeuroMab 75-173, 1:500), mouse anti-Parkin (Cell Signaling Technology 4211, 1:500) mouse anti-PINK1 mouse (NeuroMab, Clone N357/6, 1:3), anti-VDAC1 (NeuroMab 75-204 1:1000), mouse anti-ATP5a (from OXPHOS rodent antibody cocktail, Abcam ab110413, 1:2000), rabbit anti-Actin (Sigma A2066 1:1000). Primary antibodies for immunofluorescence: rat anti-GFP (Nacalai Tesque 04404-26, 1:2000). ApoTrack™ Cytochrome C Apoptosis WB Antibody Cocktail (CValpha, PDH E1alpha, cytochrome C, GAPDH) was from Abcam (1:1000, mouse), anti-Actin (1:2000, rabbit) and anti-Flag (1:1000, rabbit) were from Sigma, anti-Tom20 (1:500, rabbit) was from Santa Cruz Biotechnology. Anti-myc (9E10) and anti-HA were obtained from 9E10 and 12CA5 hybridoma lines respectively and used as supernatant at 1:100.

### Cell lines

^Flag^Parkin stably overexpressing SH-SY5Y cells are a gift from Dr. Helen Ardley (Leeds Institute of Molecular Medicine) and were previously described (Ardley et al., 2003). WT and Miro1/ Miro2 DKO mouse embryonic fibroblasts (MEFs) were characterised previously (Lopez-Domenech et al., 2018). Cells were maintained in Dulbecco’s Modified Eagle Medium (DMEM) with 4500 mg/L glucose and supplemented with Fetal Bovine Serum (FBS) (COS-7 10%; MEFs 15%), Glutamax 2 mM, Penicillin 120 μg/ml and Streptomycin 200 μg/ml, and kept at 37° C and 5% CO2 atmosphere.

### Biochemical assays

For ubiquitination assays cells were lysed in RIPA buffer (50 mM Tris pH7.4, 150 mM NaCl, 1 mM EDTA, 2 mM EGTA, 1% NP-40, 0.5% deoxycholic acid, 0.1% SDS, 1 mM PMSF and antipain, leupeptin, pepstatin at 10 μg/mL each). Lysates were then incubated on a rotating wheel at 4°C for 1 hour and then nuclei and cellular debris were spun down at 20000 g for 10 minutes. Supernatants were incubated at 4°C for 2 hours with 10 μL of a 50% slurry of anti-myc beads (Sigma).

### Primary neuronal cultures

Primary cultures were prepared from E16.5 mice essentially as in (Lopez-Domenech et al., 2016). Briefly, hippocampus from each embryo were dissected independently in ice cold HBSS. Dissected tissue was treated with 0.075% trypsin in 1mL HBSS per embryo at 37°C for 15 minutes. Tissue was washed twice and triturated to a single cell suspension. 350,000 cells were added to 6cm dishes containing 8-10 PLL-coated glass coverslips in 5mL attachment media. Media was changed to maintenance media (Neurobasal plus B27 supplement (Invitrogen)) 6-12 hours after plating. A tail section from each embryo was retained for genotyping.

### Lipofectamine transfection

Neurons were transfected at 7DIV using Lipofectamine 2000 (Invitrogen). Per 6cm dish, 1.5μg DNA was combined with 300μl Neurobasal (NB) and 2μl Lipofectamine with 300μl NB. After 5 minutes Lipofectamine was added to the DNA and incubated for 30 minutes to complex. 1.4mL of warm NB containing 0.6% glucose was added to the complex. 2mL total volume was added directly to 6cm dish and incubated for 90 minutes at 37°C. Solution was then aspirated and replaced with conditioned media. Cultures were left to rest for >72 hours prior to experimentation.

### Tissue processing

Animals of the selected age were culled by cervical dislocation or CO2 exposure. Brains were snap-frozen in liquid nitrogen and lysates produced with the appropriate lysis buffers. For histological studies animals were deeply anesthesiated and transcardially perfused with chilled 4% PFA to maintain mitochondrial integrity. Brains were further fixed by immersion in 4% PFA overnight at 4°C, cryoprotected in 30% sucrose-PBS for 24-48 hours and stored at -80°C. Tissue was serially cryosectioned in a Bright OTF-AS Cryostat (Bright Instrument, Co. Ltd.) at 30 μm thickness and stored in cryoprotective solution (30% PEG, 30% glycerol in PBS) at -20°C until used.

### Immunohistochemistry

Immunohistochemistry studies were performed in free-floating sections. Briefly, tissue sections were washed 3-5 times in PBS over 30 minutes and permeabilized in PBS-0.5% Triton-X100 over a total of 3-5 washes during 30 minutes. Sections were then blocked at RT in a solution containing 3% BSA, 10% fetal bovine serum and 0.2M glycine in PBS 0.5% Triton-X100 for 3-4 hours. Eventually, tissue was further blocked overnight at 4°C in the same blocking solution plus purified goat anti-mouse Fab-fragment (Jackson Immunoresearch) at a concentration of 50 μg/ml to reduce endogenous background. Sections were then further washed and incubated overnight at 4°C in primary antibodies prepared in blocking solution. After washing in PBS 0.5% Triton-X100 secondary antibodies were applied in blocking solution and incubated at RT for 3-4 hours. Sections were mounted on glass slides using Mowiol mounting media and stored at 4°C in the dark until documented.

### Western blotting

For western blotting 20 μg of protein from lysates of adult hippocampus or from cultured MEFs cell lines were loaded on 8-12% acrylamide gels and transferred to nitrocellulose membranes (GE Healthcare Bio-Sciences) using the Biorad system. Membranes where blocked in 4% powder skimmed milk on TBS-T for 1-2 hours. Antibodies were incubated in blocking solution (overnight at 4°C for primary antibodies or 1 hour at RT for HRP-conjugated secondary antibodies). Membranes were developed using the ECL-Plus reagent (GE Healthcare Bio-Sciences) and acquired in a chemiluminescence imager coupled to a CCD camera (ImageQuant LAS 4000mini). Densitometric analysis was performed using ImageJ software (https://imagej.nih.gov/ij/).

### Immunocytochemistry

MEF cells growing in coated coverslips or hippocampal cultures where fixed when required in 4% PFA for 10 min at RT and rinsed several times in PBS. For immunocytochemistry coverslips where permeabilized in PBS with 0.1% Triton-X100 and incubated 1 hour in blocking solution (1% BSA, 10% fetal bovine serum, 0.2M glycine in PBS with 0.1%Triton-X100). Primary antibodies where applied in blocking solution at the desired concentration and incubated for 2 hours. AlexaFluor (Invitrogen) secondary antibodies where incubated for one hour at 1:800 in blocking solution. Coverslips were mounted in Mowiol mounting media and kept in the dark at 4°C until imaged.

### Mitophagy assays

YFP-Parkin and MtDsRed expressing neurons were used for experimentation at 10-12DIV. Cultures were treated with 1uM valinomycin (Sigma) diluted in neuronal maintenance media for various timepoints ranging from 30 minutes to 5 hours prior to fixation with 4% PFA solution.

WT and Miro^DKO^ MEFs were transfected with YFP-Parkin and Miro1myc or Miro1^allR^myc when required. The next day cells were split and seeded onto fibronectin-coated coverslips at 20 μg / ml at a density of 50.000 cells / cm^2^. 24 hours later mitophagy was induced by 10 μm FCCP treatments that were applied for the selected timepoints (1, 3, 6, 12 and 24 hours). Coverslips were then fixed and stored at -20° C until processed for immunofluorescence.

### Image acquisition and analysis

#### Confocal imaging

Confocal images (1024 x 1024 pixels) were acquired on a Zeiss LSM700 upright confocal microscope (Carl Zeiss, Welwyn Garden City, UK) using a 40X or 63X oil immersion objective (NA: 1.3 and NA: 1.4 respectively). Images were processed with ImageJ software (http://imagej.nih.gov/ij/). The DeconvolutionLab2 plugin for ImageJ was used to deconvolute confocal images of primary neurons (Sage et al., 2017). Stages of Parkin induced mitophagy were scored from images taken with the 40X objective by a blinded researcher. Data collected from at least three independent experiments. For histology studies confocal images were taken using a 5X (NA: 0.16) or 10X (NA: 0.3) air objective or a 63X oil immersion objective (NA: 1.4). Acquisition parameters were kept constant over experimental conditions. All histology experiments were performed with age matched animals. For high-resolution imaging of mitoDendra expressing brains a 63X oil immersion objective (NA: 1.4) coupled to a Zeiss LSM 880 inverted confocal microscope with Airyscan technology was used. *Electron microscopy*. Brain slices were immersed and fixed with 2% paraformaldehyde, 2% glutaraldehyde, 2% sucrose in 0.1M cacodylate buffer pH7.3 and then post-fixed in 1% OsO4 / 0.1M cacodylate buffer pH7.3 at 3° C for one and a half hours. Sections were then *en bloc* stained with 0.5% uranyl acetate dH20 at 3° C for 30 min. Specimens were dehydrated in a graded ethanol-water series, infiltrated with Agar-100 resin and then hardened at 60c for 24hrs. 1 μm sections were cut and stained with 1% toluidine blue in dH20 for light microscopy. At the correct position, ultra-thin sections were cut at 70-80 nm using a diamond knife on a Reichert ultra-microtome. Sections were collected on 300 mesh copper grids and then stained with lead citrate. Sections were viewed in a Joel 1010 transition electron microscope and images recorded using a Gatan orius camera.

